# Assessing Disease Experience across the Life Span for Individuals with Osteogenesis Imperfecta: Challenges and Opportunities for Patient-Reported Outcomes (PROs) Measurement

**DOI:** 10.1101/418251

**Authors:** Laura L. Tosi, Marianne K. Floor, Christina M. Dollar, Austin P. Gillies, Members of the Brittle Bone Disease Consortium, Tracy S. Hart, David Cuthbertson, V. Reid Sutton, Jeffrey P. Krischer

## Abstract

**Background:** Patient reported outcome (PRO) information is crucial for establishing better patient-provider communication, improving shared decision making between clinicians and patients, and assessing patient responses to therapeutic interventions and increasing satisfaction with care. We used the Brittle Bones Disease Consortium (BBDC) Contact Registry for People with OI, managed by the Rare Disease Clinical Research Network (RDCRN) to (1) to evaluate the construct validity of the Patient-Reported Outcome Measurement Information System^®^ (PROMIS^®^) to record important components of the disease experience among individuals with OI; and (2) explore the feasibility of using a registry to recruit individuals with OI to report on health status. Our long-term goal is to enhance communication of health and disease management findings back to the OI community, especially those who do not have access to major OI clinical centers.

**Results:** We demonstrated the construct validity of PROMIS instruments in OI. Our results confirm that the scores from most domains differ significantly from the general US population: individuals with OI have worse symptom burden and functioning. We found no excessive floor or ceiling effects. Our study demonstrates that the BBDC Contact Registry can be used to recruit participants for online health status surveys. However, there are numerous challenges that must be addressed: lack of self-knowledge of OI type, under-representation of men, limited ethnic diversity, and imperfect questionnaire completion rates.

**Conclusion:** Our pilot study demonstrated the feasibility of using a contact registry to recruit respondents from the OI community and to obtain analyzable PROMIS data regarding disease experience. Because the results differ from the general population and avoid excessive floor and ceiling effects, PROMIS instruments can be used to assess response to therapeutic interventions in individuals with OI. Future directions will include (1) development and validation of an OI-specific patient-based classification system that aggregates persons with similar clinical characteristics and risks for complications to identify treatment needs; and (2) integrating these PRO tools into routine patient care and research studies.

## BACKGROUND

### Clinical and Epidemiologic Features of Osteogenesis Imperfecta

Osteogenesis Imperfecta (OI), commonly known as “brittle bone disease,” is a group of disorders characterized by bone fragility. OI is associated, as well, with other health problems, including scoliosis, impaired dentition, joint laxity, hearing loss, and cardiopulmonary challenges [1-16]. The severity of the disease ranges from neonatal-lethal to very mild disease with only a few bone fractures across the lifespan. Since 1979, most individuals with OI have been categorized using the Sillence phenotypic classification, which describes individuals as having mild (Type I), neonatal lethal (Type II), severely deforming (Type III), or moderately deforming (Type IV) disease [17]. However, with advances in understanding the genetic basis of OI, 18 gene-based types of OI have been proposed in the research literature, often leading to confusion on the part of patients and clinicians alike. Since the 2009 meeting of the International Nomenclature group for Constitutional Disorders ICHG of the Skeleton (INCDS), the consensus is to continue grouping the known OI syndromes into five groups based on similarities in clinical presentation. This classification preserves the primary four groups described by Sillence and adds OI type V. The different genetic causes of the OI types are recognized by listing the causative genes as subtypes of OI types I–V [18, 19]

Particularly over the past decade, there is increasing demand by the OI community to receive better information on the natural history of this disorder. Little information is available about the natural history and progression of OI in adulthood. Especially adults with OI want better delineation of health risks and evidence-based treatments. We need to know more about the patient-centered outcomes of various treatments that patients themselves report or that proxy respondents for individuals in the pediatric age ranges who cannot self-report.

### Patient-Reported Outcomes

Despite the potential for mortality and significant morbidity, no measures of patient-reported outcomes (PROs) have been designed specifically for patients with OI – certainly none with input from the OI community. Available outcome measures have been developed chiefly by medical experts, relying on physician-based assessments. Recognition has grown that individuals’ experiences and health-related quality of life (HRQOL) should be part of the assessment of new therapies and interventions. Such information is crucial for enhancing patient-provider communication, improving shared decision making between clinicians and patients (or parents), and increasing patient satisfaction with care [20-23]. Moreover, PRO data need to be linked to and analyzed with patients’ diagnostic and treatment information. This would identify clinical predictors of survivorship difficulties and, thus, facilitate early risk stratification and targeted interventions [24].

### Disease Registries and the Osteogenesis Imperfecta Patient Population

Developing data to better inform assessment and care for individuals with rare diseases is difficult; patients can be challenging to access and it can be hard to collect enough data to provide informed conclusions. One frequently recommended tool is a registry [25-31]. Registries are particularly useful in the care of individuals with a rare disorder as they can help patients gain a broader insight into their health status regardless of whether they have access to a major clinical center. The OI community has been very responsive to participating in research registries and to reporting their disease experiences. For more than a decade the OI community in the United States has embraced registry and natural history efforts sponsored by the Osteogenesis Imperfecta Foundation (OIF), a key national OI advocacy organization dedicated to supporting individuals with OI. The OIF effort began in earnest in 2005 with the establishment of the International Osteogenesis Imperfecta Registry [32]. The OIF then funded the establishment of the Linked Clinical Research Centers (LCRC) between 2009 and 2014 (five clinical sites with dedicated OI clinics); this preliminary effort demonstrated the willingness of the OI community to participate in longitudinal clinical studies, as well. [33].

Subsequently, in 2011, the OI Adult Natural History Initiative (OI ANHI) demonstrated the willingness of the OI community to participate in on-line investigations. It developed a snapshot of the health status, needs, and priorities of adults with OI. The OI ANHI web-based survey leveraged the Patient-Reported Outcome Measurement Information System^®^ (PROMIS^®^) initiative, which had been developed with support from the US National Institutes of Health since 2004 [34]. PROMIS offers a variety of methods to measure important HRQOL domains with established short forms and use of item banks for computerized-adaptive testing (CATs). The OI ANHI survey included basic demographic information, PROMIS instruments, and a detailed review of systems.

In the OI ANHI survey, PROMIS scores varied by OI disease severity (whether stratified by the 1979 Sillence classification or by patient-reported mild, moderate, or severe disease status). Scores for OI patients were often worse from those of any relevant comparison or normative population. Moreover, when OI patients were asked to rank their health concerns, such as ambulation, craniofacial and dental problems, or hearing loss (all common problems in OI), the rankings tended to differ greatly from the priorities of physicians [35]. For example, individuals with OI listed vision concerns of greater concern than cardiopulmonary disease, the opposite of the clinician perspectives.

### The Disease Experience of Persons with Osteogenesis Imperfecta

This paper reports on the latest phase of the OIF’s efforts to enhance the voice of individuals with OI, this time combining an internet-based registry with PRO measures. The OIF is part of the Brittle Bone Disorder Consortium (BBDC), an NIH funded rare-disease multi-centered project designed to provide a better basis for individuals with OI to assess their disease by tracking the natural history of OI and to support the development of other studies that further explore the disorder. Related goals are to help individuals with OI to better direct their health management and to identify areas in which new intervention strategies are needed.

### Evaluating the validity of existing PRO measures for persons with OI: A pilot

As we strive to build a standardized and robust platform to comprehensively capture the experiences of OI patients, this pilot study evaluates (1) the feasibility of using an on-line registry to recruit individuals with OI and to stratify appropriately individuals with this rare and very diverse disorder; and (2) the construct validity of PROMIS measures to record important components of the disease experience among individuals with OI. We report on these issues for an adult population of self-responders, as well as for proxy (parent) respondents for children and adolescents.

## METHODS

In order to select PRO tools with domains felt to be important to individuals with OI, we convened a diverse Steering Committee comprising individuals with OI, the parent of a child with OI, members of the Data Management and Coordinating Center, physicians specializing in OI, and representatives of the OIF. The Steering Committee reviewed all PROMIS instruments available at the time of study inception. The Steering Committee gave highest priority to using the PROMIS CAT instruments for this project. CAT is a dynamically administered computer-based test in which responses to previous completed questions, within the same PRO scale, are used to select the most appropriate next question from the PROMIS item bank. The CAT system will continue to administer questions until an ideal standard error threshold is met. Compared to fixed length questionnaires, the advantage of CAT-based assessment is reduced respondent burden which can help to improve completion rates of the questionnaire [36]. The Steering Committee selected nine PROMIS CAT-based instruments for adults covering the following HRQOL domains: Anxiety, Depression, Fatigue, Pain Behavior, Pain Interference, Physical Function, Physical Function with Mobility, Sleep Disturbance, and Satisfaction with Participation in Social Roles. For children, the Steering Committee selected PROMIS parent-proxy CAT-based instruments for patients ages 5 to 17 years covering six domains (Anxiety, Depressive Symptoms, Fatigue, Pain Interference, Mobility, and Peer Relationships). Survey respondents were expected to answer an average of 4 or 5 items per PROMIS CAT-based measure. We estimated that responding to all selected PROMIS instruments would typically take participants between 15 and 30 minutes.

PROMIS instruments were not available for children ages 2-4, therefore we also included the Pediatric Outcomes Data Collection Instrument (PODCI) which is designed to document functional status in children and adolescents with musculoskeletal disorders. We included it because of its documented validity in assessing pediatric patients with restricted mobility or disability and because of its use by the BBDC [37]. PODCI was administered using parents as the proxy respondents.

A questionnaire titled “Information About You” was developed by the study team in an effort to capture demographic and basic clinical history, as recalled by the participants. Barring geographical location and race, this information is not routinely collected during registration in the Contact Registry. The items requested in” Information about You” were the same for all participants. (Supplemental Material A)

The Rare Disease Clinical Research Network (RDCRN) Data Management and Coordinating Center, housed at the University of South Florida, manages the BBDC Contact Registry for People with OI as well as all BBDC data. The BBDC Contact Registry sent an initial registry-wide call for participation electronically on June 8, 2016, to all registry members. The registry e-mail invitation contained a link to the informed consent form for the pilot project and related questionnaires. The OIF facilitated additional recruitment, through e-mail announcements to its registered website users, social media posts, and an announcement in an electronic newsletter; all these notices encouraged interested persons to become members of the Contact Registry and enroll in the study. Once participants consented to the pilot study, they gained access to the online questionnaires/instruments. In total, the Data and Management and Coordinating Center sent 1,165 emails to 1,034 registrants, inviting them to participate in the study; 908 registrants were contacted once, 121 registrants twice, and 5 registrants three times. The data collection phase of this study closed on January 20, 2017.

To be eligible for project inclusion, respondents (including the proxies for children and adolescents) were required to be English-speaking and an adult, responding to the project either for him/herself or as a proxy for a child or adolescent age 2 to 17 years.

Because we generated a unique link to the project questionnaires for each participant (as part of the consent form), we could determine who had initiated and completed the questionnaires. We sent reminder e-mails to participants who had only partially completed the survey. Once participants completed their questionnaires, responses were downloaded to the Data Management and Coordinating Center.

Study participants entered their responses directly into online forms. All project data were collected via systems created in collaboration with the DMCC and the BBDC; these systems complied with all applicable guidelines regarding patient confidentiality and data integrity.

We conducted the entire pilot project using REDCap, a secure web-based application designed to support data capture for research studies [38]. Within its library are both the PROMIS and the PODCI. We also adapted the “Information about You” questionnaire to REDCap. The selected PROMIS, the PODCI, and the “Information about You” questionnaires were mounted on the web by the RDCRN. Participants could complete the surveys either all at once or in multiple sessions.

The order for the study survey for adults was as follows: (1) Consent; (2) Information about You; (3) PROMIS CAT Anxiety, Depression, Fatigue, Pain Behavior, Pain Interference, Physical Function or Physical Function with Mobility, Sleep Disturbance, Satisfaction with Participation in Social Roles; and 4) Comment Form. The order for children and adolescent-parent proxy survey was: (1) Consent; (2) Information about You; (3) PODCI; (4) PROMIS CAT Anxiety, Depressive Symptoms, Fatigue, Pain Interference, Mobility, Peer Relations; and (5) Comment Form.

At the completion of the data collection phase, participant data were downloaded to the Data Management and Coordinating Center and analyzed using SAS version 9.4 (SAS Institute, Cary, NC) and R software version 3.4.2 [39]

We scored and standardized responses on all PROMIS instruments. Each participant’s pattern of responses was converted into a standardized T-score, based on the U.S. general population for adults (and a mix of general and clinical populations for children), with a mean of 50 and a standard deviation (SD) of 10. The standardized T-score is reported as the final score for each participant. To determine whether the results of the PROMIS instruments differed from those for the general US population, we conducted one-sample t-tests on normally distributed results and one-sample Wilcoxon Rank tests on non-normally distributed data. Floor and ceiling effects in the PROMIS^®^ CAT instruments were assessed by measuring the proportion of subjects who scored all the questions for an instrument at either the lowest or the highest possible score range. Higher scores on these instruments are indicative of greater disease burden.

The PODCI was scored according to the Version 2.0 Scoring Algorithm from the AAOS [40]. We input the collected scores for the instrument into the scoring algorithm which produced mean, standardized, and normalized scores for each domain. These were then compared to PROMIS domains using Pearson’s correlation test to assess for congruent validity (a form of construct validity).

IRB approval for the pilot project was provided by the University of South Florida, the home institution of the Contact Registry.

## RESULTS

Three hundred individuals with self-reported OI or serving as parent proxies, representing a wide range of self-reported disease severity, enrolled in the study. Of the 300 original enrollees, 290 (97%) individuals filled out some portion of the survey questionnaires; 10 individuals consented but did not start any questionnaire. Of the 290 respondents, 27 were parent-proxies for children ages 2-4 years, 65 were parent-proxies for children ages 5-17 years, and 198 were adults with OI of ages 18+ years. In all, 273 of the 290 individuals or their proxy respondents, opened all designated questionnaires (94%); 17 opened some questionnaires.

Higher completion rates were noted for adult-specific instruments than for child-specific instruments; 94-98% of adults completed all nine adult-specific PROMIS instruments, whereas 79-86% of parent-proxies completed all six child-specific PROMIS instruments (p<0.01). Similarly for the six individual PODCI domains we noted lower parent-proxy completion rates ranging from 76-90%. (Table 1). Only 64/92 (70%) parent proxies completed all PODCI domains.

**Table 1:** Questionnaire Completion Table

| **Total Instrument Completion** |  |  |  |
| --- | --- | --- | --- |
| **Instrument** | **Completed Surveys** | **Percent** | **Total N** |
| Information About You | 290 (+/- specific items) | 100% | 290 |
| PODCI Parent Reported (1/6 Domain Completed)* | 84 | 91.3% | 92 |
| Upper Extremity | 74 | 80.4% | 92 |
| Transfer Mobility | 79 | 85.9% | 92 |
| Sports Physical Function | 75 | 81.5% | 92 |
| Pain Comfort | 83 | 90.2% | 92 |
| Happiness | 70 | 76.1% | 92 |
| Global Function | 70 | 76.1% | 92 |
| PODCI Parent Reported (6/6 Domains Completed) | 64 | 69.6% | 92 |
| Proxy-Reported Anxiety Survey | 56 | 86.2% | 65 |
| Proxy-Reported Depressive Symptoms Survey | 54 | 83.1% | 65 |
| Proxy-Reported Fatigue Survey | 52 | 80.0% | 65 |
| Proxy-Reported Pain Interference Survey | 51 | 78.5% | 65 |
| Proxy-Reported Mobility Survey | 52 | 80.0% | 65 |
| Proxy-Reported Peer Relationships Survey | 52 | 80.0% | 65 |
| Adult Anxiety Survey | 193 | 97.5% | 197 |
| Adult Depression Survey | 192 | 97.0% | 197 |
| Adult Fatigue Survey | 190 | 96.0% | 197 |
| Adult Pain Behavior Survey | 190 | 96.0% | 197 |
| Adult Pain Interference Survey | 188 | 95.0% | 197 |
| Adult Physical Function Survey | 102 | 94.4 | 197 |
| Adult Physical Function with Mobility Aid** | 85 |  |  |
| Adult Satisfaction with Participation in Social Roles | 186 | 93.9% | 197 |
| Adult Sleep Disturbance Survey | 188 | 95.0% | 197 |
| Comment Form | 252 | 86.9% | 290 |

\*Does not include one 18 year old, and does not delineate between 2-4 and 5-17
\*\*Used in lieu of PROMIS® Adult CAT Physical Function instrument if the participant states the use of a mobility aid

The majority of PROMIS instruments required, on average, participants to answer fewer than six questions to enable us to calculate a PRO score (mean = 5.53 questions). The domain that required the most questions was Adult Physical Function with Mobility Aid (mean = 9.39 questions).

General demographics and self-reported clinical history of both adult and child subjects came from the “Information about You” question set. Responses are summarized in Table 2.

**Table 2:**
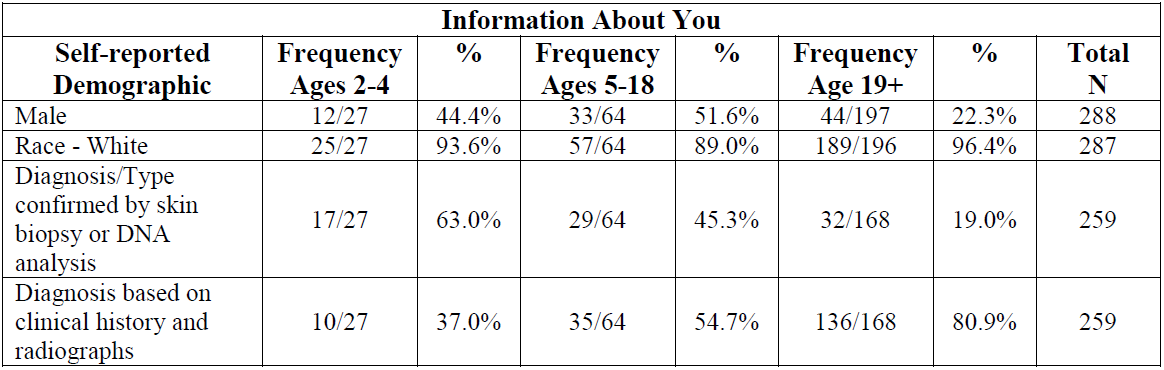
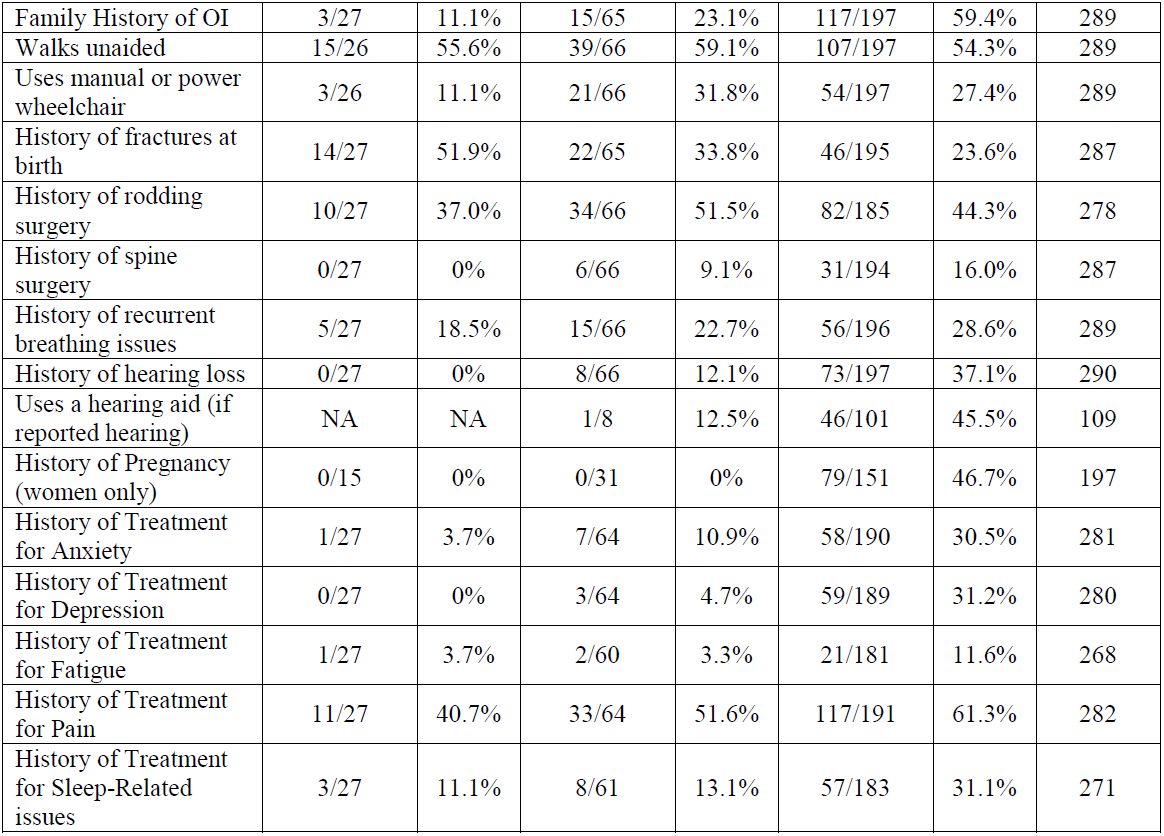
Information About You

### Age and Sex

Figure 1 displays the age ranges of respondents, stratified by sex. In the adult population, 78% (154/198) were female; in the pediatric population, 51% (46/90) were female. The proportion of male vs female subjects was essentially balanced in children and adolescents, while female respondents far out-numbered males respondents in adults. Despite the lower response by males, there were at least some male respondents in most adult age cohorts.

**Figure 1:**
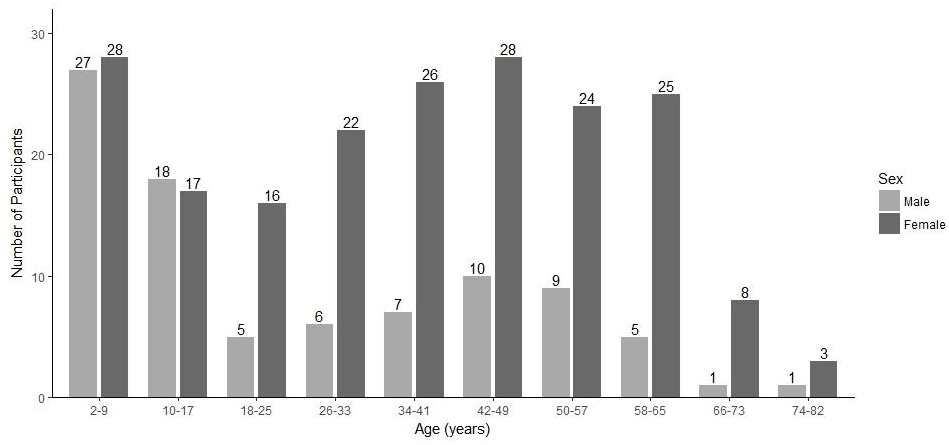
Distribution of Patients by Age and Sex

### Race and Ethnicity

Just under 95% (271/287) of respondents self-identified as white; (8% (23/287) as Hispanic, Latino, or Spanish Origin); 3% (8/287) as Asian; 1% (3/287) as Black; and 1% (3/287) as American Indian or Alaskan Native. One person declined to report race and one person listed race as Unknown. Of note, 77% (231/300) of enrollees were located in the U.S. and 23% (23/300) were located abroad. The demographic data of respondents who participated in this pilot study mirrored those in the Contact Registry except that 5% of Registry members self-reported as being of multiple races.

### OI Type

Figure 2 depicts the self-reported distribution of OI type within our study cohort. Of all 290 participants, 20% did not know their OI type. Participants spanned the spectrum of common self-reported OI types. Four patients reported that they were Type II (perinatal lethal form); of these, all were adults and three were older than 50. The mean age (40.2 years) of those who did not know their OI Type was 8.59 years older than those who said they knew their OI Type (31.42 years). The difference is statistically significant (*p*<0.01)

**Figure 2:**
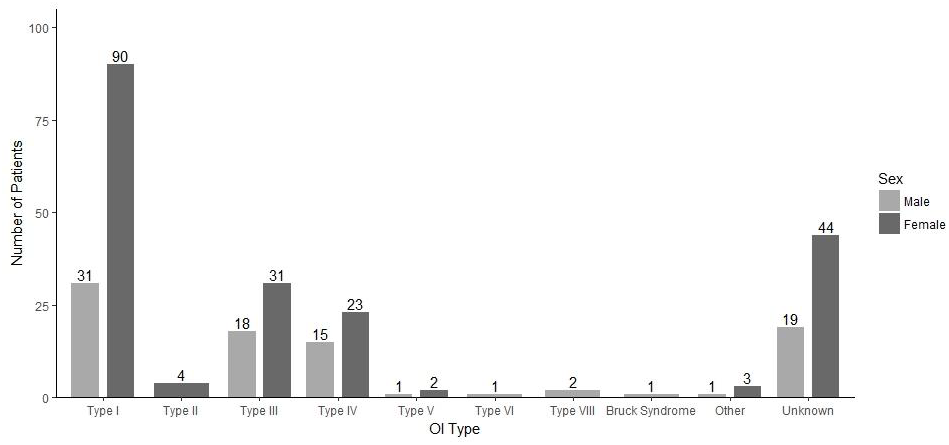
Distribution of Patients by Age and OI Type.

### OI Diagnosis

Of those patients (N = 78) who reported to know how they had been diagnosed, 12% (30/159) reported that clinicians made the diagnosis through skin biopsy or collagen studies and 19% (48/159) reported was via blood or DNA studies. 70% reported that the diagnosis was based on “clinical history and radiographs highly suggestive of Osteogenesis Imperfecta.”

T-scores for all PROMIS instruments showed a reasonable mean and range. All PROMIS domains for adults showed a statistically significant difference between OI patients and the general population, as did four parent-proxy instruments: Anxiety, Pain Interference, Mobility and Sleep Disturbance in children/adolescents (Table 3).

**Table 3:**
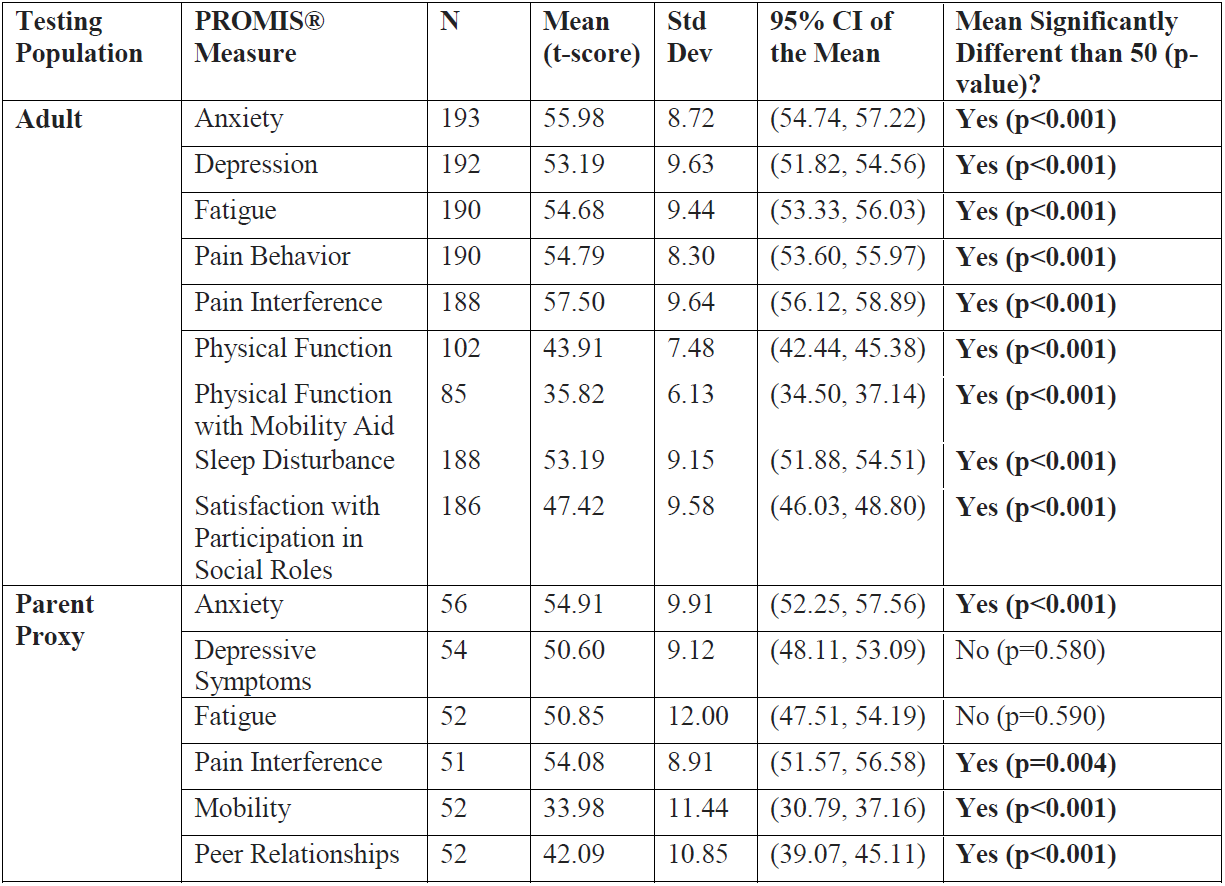
PROMIS Instrument Analysis

| Testing Population | PROMIS® Measure | N | Mean (t-score) | Std Dev | 95% CI of the Mean | Mean Significantly Different than 50 (p-value)? |
| --- | --- | --- | --- | --- | --- | --- |
| **Adult** | Anxiety | 193 | 55.98 | 8.72 | (54.74, 57.22) | **Yes (p<0.001)** |
|  | Depression | 192 | 53.19 | 9.63 | (51.82, 54.56) | **Yes (p<0.001)** |
|  | Fatigue | 190 | 54.68 | 9.44 | (53.33, 56.03) | **Yes (p<0.001)** |
|  | Pain Behavior | 190 | 54.79 | 8.30 | (53.60, 55.97) | **Yes (p<0.001)** |
|  | Pain Interference | 188 | 57.50 | 9.64 | (56.12, 58.89) | **Yes (p<0.001)** |
|  | Physical Function | 102 | 43.91 | 7.48 | (42.44, 45.38) | **Yes (p<0.001)** |
|  | Physical Function with Mobility Aid | 85 | 35.82 | 6.13 | (34.50, 37.14) | **Yes (p<0.001)** |
|  | Sleep Disturbance | 188 | 53.19 | 9.15 | (51.88, 54.51) | **Yes (p<0.001)** |
|  | Satisfaction with Participation in Social Roles | 186 | 47.42 | 9.58 | (46.03, 48.80) | **Yes (p<0.001)** |
| **Parent Proxy** | Anxiety | 56 | 54.91 | 9.91 | (52.25, 57.56) | **Yes (p<0.001)** |
|  | Depressive Symptoms | 54 | 50.60 | 9.12 | (48.11, 53.09) | No (p=0.580) |
|  | Fatigue | 52 | 50.85 | 12.00 | (47.51, 54.19) | No (p=0.590) |
|  | Pain Interference | 51 | 54.08 | 8.91 | (51.57, 56.58) | **Yes (p=0.004)** |
|  | Mobility | 52 | 33.98 | 11.44 | (30.79, 37.16) | **Yes (p<0.001)** |
|  | Peer Relationships | 52 | 42.09 | 10.85 | (39.07, 45.11) | **Yes (p<0.001)** |

For adults, Pain Behavior and Pain Interference had floor effects above 10% as did Fatigue, Pain Interference and Mobility in children. All were under the generally accepted 15% cut-off point. No important ceiling effects were detected in adult or parent-proxy instruments. (Table 4).

**Table 4:**
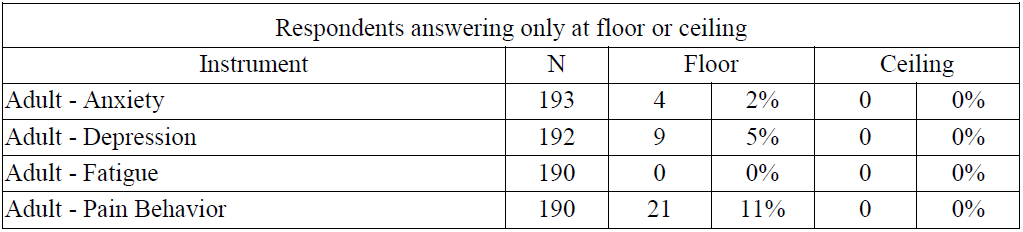
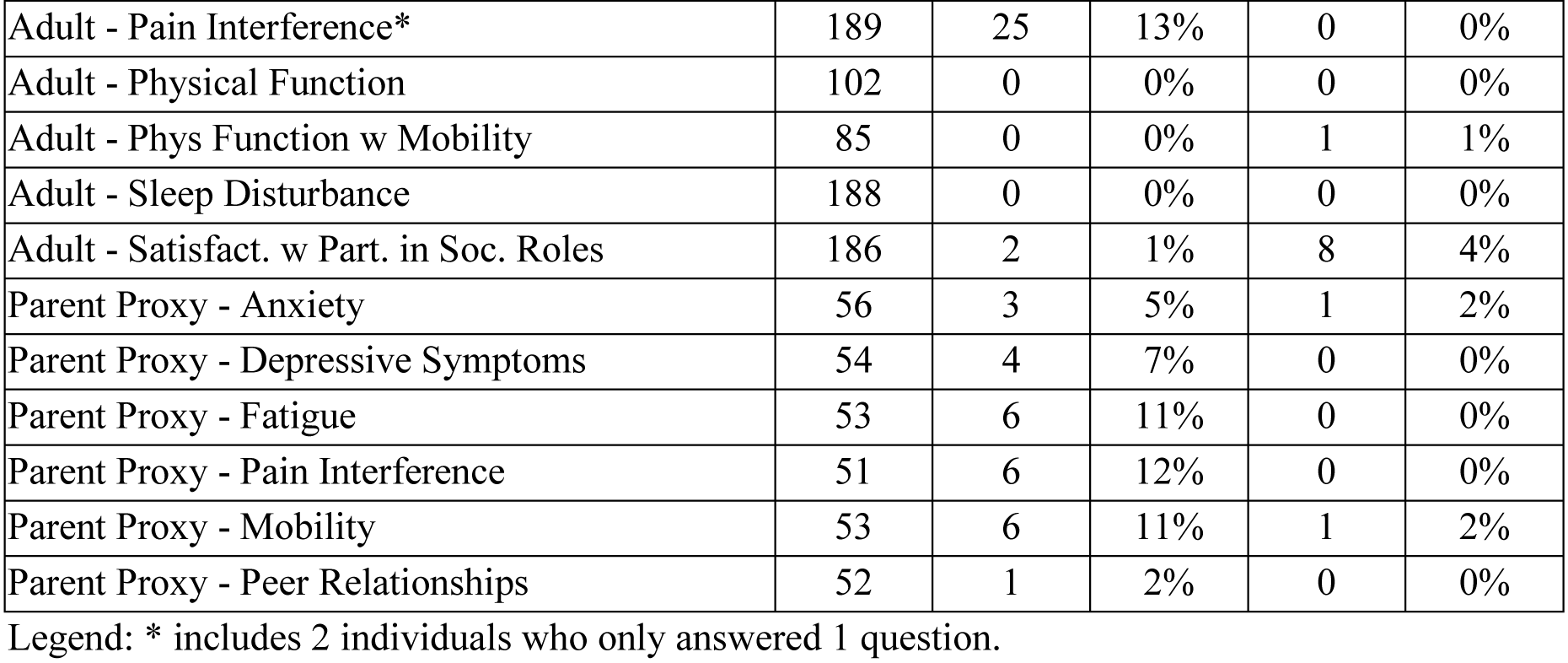
Floor and Ceiling Effects

In addition, we compared the questionnaire results for individuals with OI Type I with the results for all other types combined, excluding individuals with unknown types, to determine whether PROMIS instruments can distinguish among OI Type I and the other, more severe, types (evaluation of known-group validity). Two adult and one parent-proxy PROMIS instruments were able to discriminate between OI Type I and self-reported OI Types III and IV grouped together (*p*<0.01): Adult Physical Function (with or without Mobility Aid), Adult Sleep Disturbance, and Parent Proxy Mobility.

Of the 65 parent proxy respondents for children ages 5-17, 55 filled out both the PODCI and at least one PROMIS instrument. All six parent proxy PROMIS instruments showed statistically significant correlations with at least one PODCI element, with the strongest correlations seen between those instruments focused on physical function, which provides evidence for convergent validity of PROMIS measures. Several outcomes are essentially opposites (e.g. happiness vs anxiety), thus those correlations are negative. (Table 5)

**Table 5:**
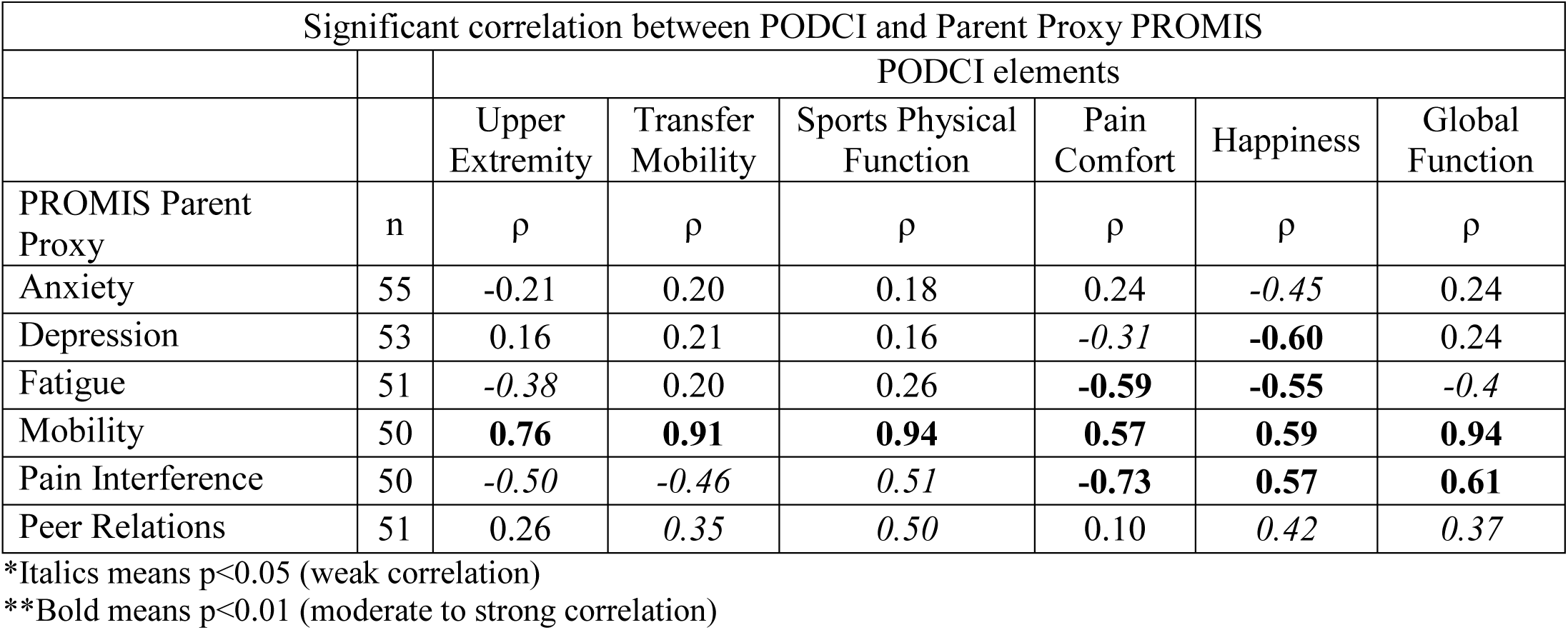
PODCI and PROMIS correlation

| Significant correlation between PODCI and Parent Proxy PROMIS |  |  |  |  |  |  |  |
| --- | --- | --- | --- | --- | --- | --- | --- |
|  |  | PODCI elements |  |  |  |  |  |
|  |  | Upper Extremity | Transfer Mobility | Sports Physical Function | Pain Comfort | Happiness | Global Function |
| PROMIS Parent Proxy | n | $\rho$ | $\rho$ | $\rho$ | $\rho$ | $\rho$ | $\rho$ |
| Anxiety | 55 | -0.21 | 0.20 | 0.18 | 0.24 | *-0.45* | 0.24 |
| Depression | 53 | 0.16 | 0.21 | 0.16 | *-0.31* | **-0.60** | 0.24 |
| Fatigue | 51 | *-0.38* | 0.20 | 0.26 | **-0.59** | **-0.55** | *-0.4* |
| Mobility | 50 | **0.76** | **0.91** | **0.94** | **0.57** | **0.59** | **0.94** |
| Pain Interference | 50 | *-0.50* | *-0.46* | *0.51* | **-0.73** | **0.57** | **0.61** |
| Peer Relations | 51 | 0.26 | *0.35* | *0.50* | 0.10 | *0.42* | *0.37* |

\*Italics means $p < 0.05$ (weak correlation)
\*\*Bold means $p < 0.01$ (moderate to strong correlation)

## DISCUSSION

Our pilot study provides evidence for the feasibility and construct validity of using PROMIS instruments to record important components of the disease experience among individuals with OI. The quantitative results from both PROMIS and PODCI can provide clinicians and researchers with a yardstick for assessing overall need for treatment and determining the success (or failure) of an intervention. In addition, our results confirm that the scores from most domains differ significantly from the general US population: individuals with OI have higher disease burden and worse functioning.

There were two exceptions of “no differences” from the general population: the parent-proxy assessments of depression and fatigue. At least two reasons could explain this finding. One is that the reference population for the PROMIS Pediatric and the PROMIS Proxy measures is a mix of both representatives of the US general population and representatives of children with other clinical conditions or disorders. Thus, it may be that the children with OI are similar to this mixed clinical/general population on these two PROs. The second reason may be that parents have more difficulty in evaluating their child’s depression and fatigue levels since these are unobservable symptoms (as opposed to more observable outcomes such as mobility).

We found few ceiling effects. Five instruments demonstrated floor effects between 11% and 15% which is in acceptable range given that floor effects for symptom measures reflect the patient is not experiencing the symptom, such as pain.

For children and adolescents, all six parent proxy PROMIS instruments showed some correlation with at least one PODCI element, providing evidence for convergent validity. The Parent Proxy – Mobility PROMIS instrument correlated strongly with the PODCI which has been used in the LCRC, the BBDC, as well as this project. The results of the correlation analysis between the PODCI and six PROMIS^®^ instruments speak to the potential value of expanding the use of PROMIS instruments for children and adolescents in the BBDC. Although the PODCI and PROMIS instruments are not measuring precisely the same aspects of HRQOL, and although the number of relevant respondents in this pilot study was relatively small, we judge that these results support the validity of PROMIS CAT instruments in measuring disease severity in children and adolescents for OI and that use of PROMIS CAT may possibly lead to higher completion rates of on-line questionnaires in future studies because they ask fewer questions of participants than fixed-length questionnaires like PODCI.

PROMIS CATs were not able to differentiate individuals by OI severity. Patients with OI and clinicians often disagree on the level of disease burden experienced. Anecdotal evidence suggests that patients with milder types of OI may have different expectations for quality of life than patients with more affected or severe types. For example, the young man with mild OI may feel severely burdened because he cannot play football with his peers and/or the individual with severe III OI may feel extremely mobile and unrestricted because they have acquired a top-of-the-line wheelchair which allows them to participate in school, family, job, and social activities easily. There is an extensive literature that illustrates that, across numerous chronic diseases, people adapt to their situation or condition, and then, within that context have expectations for quality of life that might seem surprising to people not suffering from that condition [41, 42]. It is also possible that since OI type was self-reported in this study, incorrect identification of OI type is contributing to these findings.

In addition, our pilot study demonstrates that the BBDC Contact Registry for People with OI can be used to recruit participants for online surveys regarding health status. This capability will allow researchers to better understand the OI community’s perspective on health status and quality of life; it will also permit users to capture quantitative PROs using tools such as PROMIS^®^ or PODCI to explore domains that patients identify as important. However, it also underscores the importance of challenges to using registries that must be addressed to expand our understanding of the health issues faced by individuals with OI and then communicate health and disease management findings back to the OI community, especially those who do not have access to major OI clinical centers.

### Self-knowledge of OI Type

A significant proportion of respondents did not know their OI type. This was particularly prevalent in older individuals, a troubling problem as we seek to expand our knowledge of the natural history of OI in adulthood. We believe this result reflects confusion regarding the current and historical classification of OI. Prior to the publication of the 1979 Sillence classification of OI, patients were typically described as having “congenita” or tarda” (fractures and bowing occurring after birth) disease. This has led many older patients to describe themselves as having type 1 or 2 disease. Although 18 OI types have been described in the research literature, Sillence and others recently pleaded to simplify the classification of OI to mild, moderate, and severe. The rationale is that OI type is artificial and sometimes not helpful. They contend that this simplified categorization, focusing on broad clinical findings, particularly clinical and historical data, fracture frequency, bone densitometry, level of mobility, and patient report tools [19, 43-47], can be used to enhance communication between patients and professionals.

We believe that a critical next step for the BBDC research team is leveraging the clinical data of the BBDC Natural History Study to validate a PRO question set that establishes whether patients have mild, moderate, or severe disease. Patient reports can be validated against known clinical data as well as their clinicians’ perception of disease severity. Broad dissemination of this question set and inclusion of its contents at the beginning of all surveys and studies will standardize data collection, and, most important, help members of the OI community have a better understanding of which study results are relevant to them.

Moving forward, PRO results will be particularly helpful if they demonstrate an association between patients’ perspectives of their own disease experience and objective clinical data stratified by OI type. To date, only a limited number of PROMIS instruments address physiologic symptoms, such as “breathlessness”. However, new instruments are becoming available. Where there are no PROMIS measures, there must be a decision either to develop a new measure (time consuming and costly) or use another PRO measure available in the field that yields good psychometric properties.

### Recruitment

The Contact Registry succeeded in attracting a meaningful number of patients with OI who were willing to complete an extensive set of questionnaires. However, while the demographic data on those enrolled in the registry closely approximated participation in this study, neither the Registry nor the study cohorts mirror what is known about the epidemiology of the disorder in the community. Participants in this pilot study were far less diverse in terms of sex and race/ethnicity than the known OI community. Expanded recruitment efforts will clearly be needed for future studies.

Similarly, the international prevalence of persons with OI is estimated to be between 1 and 2 per 20,000 individuals. For the United States, this translates to more than 32,000 possible respondents, yet, at the time we solicited participants in this pilot study, we had only attracted approximately 1,000 US subjects to join the BBDC Contact Registry [48]. The OIF has already taken note of this finding and is redoubling its efforts to increase enrollment. Of note, 23% of enrollees in this pilot study were from outside the United States, which speaks to the need to ensure that research findings can be communicated back to the broad OI community.

The original call to participate in this pilot study recruited 100 individuals in a day, but another 2 months were needed to fulfil the recruitment goal of 300. This suggests that although a small cohort of individuals is eager to participate in research, this enthusiasm may not extend to the entire OI community. Moreover, participants were limited to English speakers and those who had access to computers. Some members of the community may have been prevented from participating because of disease limitations or personal time constraints.

In the same vein, we had planned to link the data gathered by this pilot study to existing clinical data collected through the BBDC, so that we could associate the range of clinical findings found on examination with the range of T-scores noted in PROMIS-assessed PROs. However, only 21 of the respondents are known to have participated in the BBDC and had data available for linkage and analysis. Although the registry is a component of the BBDC, we did not specifically recruit individuals who are participating in the BBDC OI Longitudinal Study for this study. One plausible explanation for this shortfall is that those already involved in the Longitudinal Study may believe that they are sufficiently involved and making an adequate contribution to the knowledge base about OI.

### The Match (or Mismatch) between Self- and Proxy-Responses for Children

Parent-proxy reported measures of PRO results may well differ from child self-reported results, especially for adolescents and possibly for older children [49]. For this study, we did not ask children to complete self-reported instruments; we relied on parent-proxy instruments. This prevented analysis of congruency between the self-perceived disease experiences of children and the estimations of caretakers. Future studies will need to explore the similarities and differences found in parent-proxy and patient PRO responses, at least for the pediatric age groups for which reliable and valid self-report measures exist. Including tools that can be answered by older children and adolescents is essential.

### Respondent Burden

CATs instruments were developed to reduce respondent burden by limiting the number of questions that participants need to answer, and our results underscore the importance of that philosophy [36]. The PROMIS CATs for this study usually required between about four and six answers for any given domain. Indeed, in examining the 15 PROMIS instruments alone, the average number of questions answered per individual, per instrument was generally fewer than six questions. Across all the self-report (or proxy-report) domains in PROMIS, adults probably had to respond to 40-50 items. Understandably, completion rates for PODCI were lower. It required response to up to 88 questions (covering 6 domains). Adult participants were not required to complete any instrument as burdensome as the PODCI.

Data regarding the average time needed to administer our study might have enhanced our understanding of study completion rates. Unfortunately, our platform was not able to time participants as they completed the survey instruments. Thus, we cannot know how much faster, on average, participants might be able to complete a PROMIS^®^ instrument than the PODCI.

### Recognizing the voice of the patient

Finally, but perhaps most important, the issues of concern to patients with OI frequently differ from the typical clinician focus. Indeed, the PROMIS instruments chosen for the study are not typically topics covered in a routine clinic visit. The wide range of existing PROMIS instruments offers an opportunity to explore a variety of health concerns for which there may not be time to discuss during a standard clinic visit.

## Conclusion

Our pilot study demonstrated the feasibility of using the RDCRN BBDC Contact Registry for People with OI to recruit respondents from the OI community and to obtain analyzable PROMIS^®^ and PODCI data regarding their disease experience. The tools selected performed well, demonstrating statistically significant scores relative to the general population and no excessive floor or ceiling effects. We will therefore be incorporating PRO instruments into our BBDC longitudinal study to assess health status, outcomes and responses to interventions in individuals with OI. Significant work remains ahead, however, if we are to meet our broader goal of improving quality of life for individuals with OI by improving their access to OI-specific health information, regardless of whether they have access to a major OI clinical center.

Our next step will be to leverage the intimate knowledge of OI that persons with this rare, extremely heterogeneous disorder hold *and* the clinical data accumulated by the BBDC Natural History Study. Doing so will allow us to validate an OI-specific patient-based classification system that aggregates persons with similar clinical characteristics and risks for complications as a basis for identifying treatment needs and developing disease management recommendations as well as generating hypotheses for pharmaceutical and clinical management studies. We will also expand our identification of existing PRO measures (e.g., PROMIS) and/or develop new PRO measures in partnership with the OI community to detect discernable differences in health status as a basis for evaluating health status and treatment results.

## Abbreviations

BBDC: Brittle Bones Disease Consortium HRQOL: health-related quality of life
OI: Osteogenesis Imperfecta
OIF: Osteogenesis Imperfecta Foundation
PRO: patient reported outcome
PROMIS^®^: Patient-Reported Outcome Measurement Information System^®^ RDCRN Rare Disease Clinical Research Network
PODCI: Pediatric Outcomes Data Collection Instrument
INCDS: International Nomenclature group for Constitutional Disorders ICHG of the Skeleton
LCRC: Linked Clinical Research Centers
OI ANHI: OI Adult Natural History Initiative
CAT: Computerized-adaptive technology

## Declarations

### •Consent for publication

Not applicable.

### •Availability of data and material

The datasets generated and/or analysed are not currently publicly available. They will be available via dbGaP one year after publication of planned analyses.

### Authors’ contributions

LLT: Study design, statistical analysis, manuscript preparation, final approval MKF: Contribution: Study design, statistical analysis, manuscript preparation, final approval

CMD: Contribution: Statistical analysis, manuscript preparation, final approval

APG: Contribution: Statistical analysis, manuscript preparation, final approval

TSH: Contribution: Study design, manuscript preparation, final approval

DC: Contribution: Study design, performed measurements, statistical analysis, manuscript preparation, final approval

VRS: Contribution: Study design, manuscript preparation, final approval

JPK: Contribution: Study design, performed measurements, statistical analysis, manuscript preparation, final approval

## Acknowledgements

We wish to thank the members of the OI community who served on our Steering Committee: Michelle Burka, Lauren Brown, Donald Gardner, Barbara Simmonds, and Susan Wilson. We also wish to thank Kathleen N. Lohr and Bryce Reeve for their editorial suggestions.

## Appendix A Information About You Questionnaire

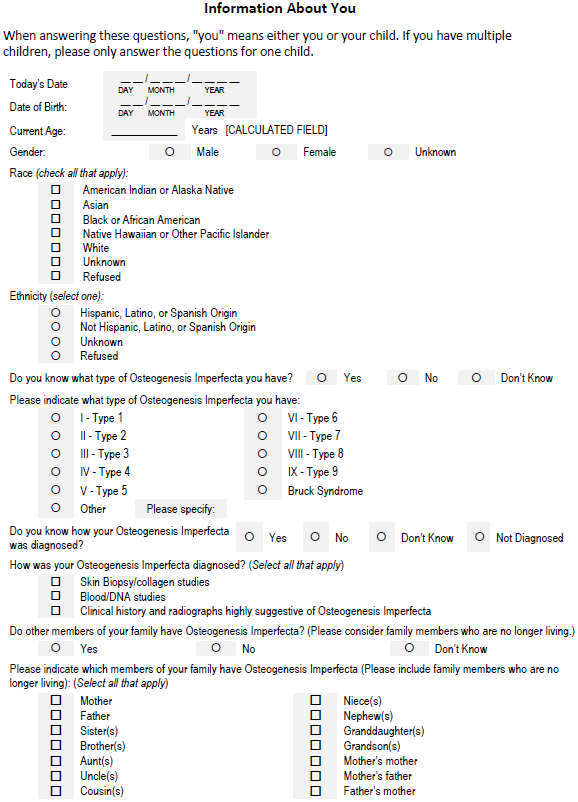
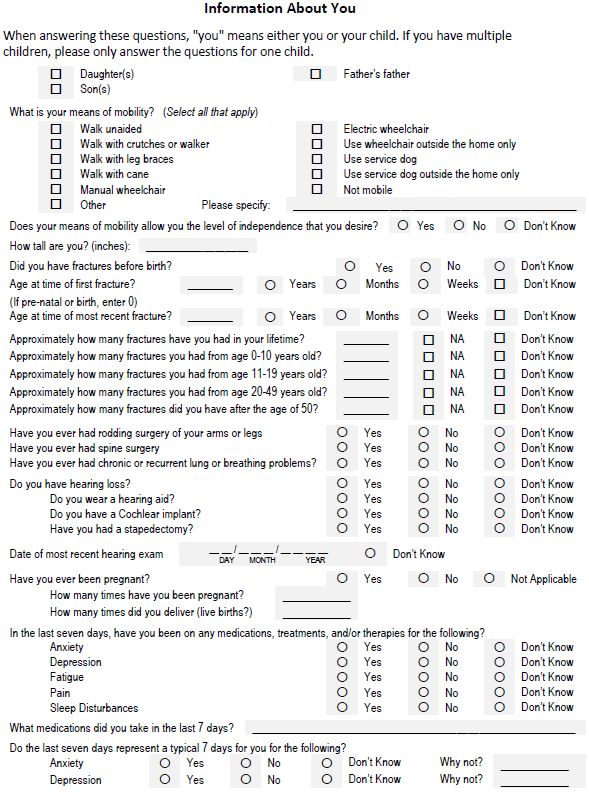
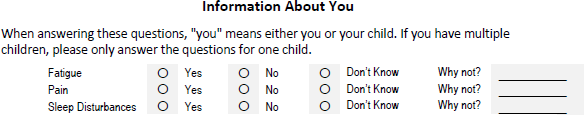

## REFERENCES

1. Augustin G, Jelincic Z, Majerovic M, Stefancic L. Carcinoma of left colon presenting as mechanical obstruction in a patient with osteogenesis imperfecta type III. J Inherit Metab Dis. 2007; 30:109–110.

2. Basel D, Steiner RD. Osteogenesis imperfecta: recent findings shed new light on this once well-understood condition. Genet Med. 2009; 11:375–385.

3. Bilkay U, Tiftikcioglu YO, Mezili C. Management of nasal deformity in osteogenesis imperfecta. J Craniofac Surg. 2010; 21:1465–1467.

4. Bonita RE, Cohen IS, Berko BA. Valvular heart disease in osteogenesis imperfecta: presentation of a case and review of the literature. Echocardiography. 2010; 27:69–73.

5. Byra P, Chillag S, Petit S. Osteogenesis imperfecta and aortic dissection. Am J Med Sci. 2008; 336:70–72.

6. Hansen B, Jemec GB. The mechanical properties of skin in osteogenesis imperfecta. Arch Dermatol. 2002; 138:909–911.

7. Li HY, Fang TJ, Lin JL, Lee ZL, Lee LA. Laryngomalacia causing sleep apnea in an osteogenesis imperfecta patient. Am J Otolaryngol. 2002; 23:378–381.

8. Litos M, Michala S, Brown R. Osteogenesis imperfecta and pregnancy. Eur J Obstet Gynecol Reprod Biol. 2008; 136:126–127.

9. McKiernan FE. Musculoskeletal manifestations of mild osteogenesis imperfecta in the adult. Osteoporos Int. 2005; 16:1698–1702.

10. Petruzzellis M, De Blasi R, Lucivero V, Sancilio M, Prontera M, Tinelli A, Mezzapesa DM, Federico F. Cerebral aneurysms in a patient with osteogenesis imperfecta and exon 28 polymorphism of COL1A2. AJNR Am J Neuroradiol. 2007; 28:397–398.

11. Pillion JP, Shapiro J. Audiological findings in osteogenesis imperfecta. J Am Acad Audiol. 2008; 19:595–601.

12. Radunovic Z, Wekre LL, Diep LM, Steine K. Cardiovascular abnormalities in adults with osteogenesis imperfecta. Am Heart J. 2011; 161:523–529.

13. Saeves R, Lande Wekre L, Ambjornsen E, Axelsson S, Nordgarden H, Storhaug K. Oral findings in adults with osteogenesis imperfecta. Spec Care Dentist. 2009; 29:102–108.

14. Scott A, Kashani S, Towler HM. Progressive myopia due to posterior staphyloma in Type I Osteogenesis Imperfecta. Int Ophthalmol. 2005; 26:167–169.

15. Takahashi S, Okada K, Nagasawa H, Shimada Y, Sakamoto H, Itoi E. Osteosarcoma occurring in osteogenesis imperfecta. Virchows Arch. 2004; 444:454–458.

16. Arponen H, Makitie O, Haukka J, Ranta H, Ekholm M, Mayranpaa MK, Kaitila I, Waltimo-Siren J. Prevalence and natural course of craniocervical junction anomalies during growth in patients with osteogenesis imperfecta. J Bone Miner Res. 2012; 27:1142–1149.

17. Sillence DO, Senn A, Danks DM. Genetic heterogeneity in osteogenesis imperfecta. J Med Genet. 1979; 16:101–116.

18. Warman ML, Cormier-Daire V, Hall C, Krakow D, Lachman R, LeMerrer M, Mortier G, Mundlos S, Nishimura G, Rimoin DL, et al. Nosology and classification of genetic skeletal disorders: 2010 revision. Am J Med Genet A. 2011; 155A:943–968.

19. Bonafe L, Cormier-Daire V, Hall C, Lachman R, Mortier G, Mundlos S, Nishimura G, Sangiorgi L, Savarirayan R, Sillence D, et al. Nosology and classification of genetic skeletal disorders: 2015 revision. Am J Med Genet A. 2015; 167A:2869–2892.

20. Calvert M, Kyte D, Mercieca-Bebber R, Slade A, Chan AW, King MT, Hunn A, Bottomley A, Regnault A, Ells C, et al. Guidelines for Inclusion of Patient-Reported Outcomes in Clinical Trial Protocols: The SPIRIT-PRO Extension. JAMA. 2018; 319:483–494.

21. Weldring T, Smith SM. Patient-Reported Outcomes (PROs) and Patient-Reported Outcome Measures (PROMs). Health Serv Insights. 2013; 6:61–68.

22. Kotronoulas G, Kearney N, Maguire R, Harrow A, Di Domenico D, Croy S, MacGillivray S. What is the value of the routine use of patient-reported outcome measures toward improvement of patient outcomes, processes of care, and health service outcomes in cancer care? A systematic review of controlled trials. J Clin Oncol. 2014; 32:1480–1501.

23. Bresnahan BW, Rundell SD. Including patient-reported outcomes and patient-reported resource-use questionnaires in studies. Acad Radiol. 2014; 21:1129–1137.

24. Snyder CF, Jensen RE, Segal JB, Wu AW. Patient-reported outcomes (PROs): putting the patient perspective in patient-centered outcomes research. Med Care. 2013; 51:S73–79.

25. Dy CJ, Bumpass DB, Makhni EC, Bozic KJ. The Evolving Role of Clinical Registries: Existing Practices and Opportunities for Orthopaedic Surgeons. J Bone Joint Surg Am. 2016; 98:e7.

26. Steinhagen-Thiessen E, Stroes E, Soran H, Johnson C, Moulin P, Iotti G, Zibellini M, Ossenkoppele B, Dippel M, Averna MR. The role of registries in rare genetic lipid disorders: Review and introduction of the first global registry in lipoprotein lipase deficiency. Atherosclerosis. 2017; 262:146–153.

27. Richesson RL, Young K, Lloyd J, Adams T, Guillette H, Malloy J, Krischer JP. An automated communication system in a Contact Registry for persons with rare diseases: tools for retaining potential clinical research participants. AMIA Annu Symp Proc. 2007:1094.

28. Richesson RL, Lee HS, Cuthbertson D, Lloyd J, Young K, Krischer JP. An automated communication system in a contact registry for persons with rare diseases: scalable tools for identifying and recruiting clinical research participants. Contemp Clin Trials. 2009; 30:55–62.

29. Richesson RL, Sutphen R, Shereff D, Krischer JP. The Rare Diseases Clinical Research Network Contact Registry update: features and functionality. Contemp Clin Trials. 2012; 33:647–656.

30. Krischer JP, Gopal-Srivastava R, Groft SC, Eckstein DJ. The Rare Diseases Clinical Research Network’s organization and approach to observational research and health outcomes research. J Gen Intern Med. 2014; 29 Suppl 3:S739–744.

31. Yimgang DP, Brizola E, Shapiro JR. Health outcomes of neonates with osteogenesis imperfecta: a cross-sectional study. J Matern Fetal Neonatal Med. 2016; 29:3889–3893.

32. Yimgang DP, Shapiro JR. Pregnancy outcomes in women with osteogenesis imperfecta. J Matern Fetal Neonatal Med. 2016; 29:2358–2362.

33. Patel RM, Nagamani SC, Cuthbertson D, Campeau PM, Krischer JP, Shapiro JR, Steiner RD, Smith PA, Bober MB, Byers PH, et al. A cross-sectional multicenter study of osteogenesis imperfecta in North America - results from the linked clinical research centers. Clin Genet. 2015; 87:133–140.

34. Health Measures: Transforming How Health is Measured. [http://www.healthmeasures.net/explore-measurement-systems/promis] Accessed 8 April 2018.

35. Tosi LL, Oetgen ME, Floor MK, Huber MB, Kennelly AM, McCarter RJ, Rak MF, Simmonds BJ, Simpson MD, Tucker CA, McKiernan FE. Initial report of the osteogenesis imperfecta adult natural history initiative. Orphanet J Rare Dis. 2015; 10:146.

36. Djaja N, Janda M, Olsen CM, Whiteman DC, Chien TW. Estimating Skin Cancer Risk: Evaluating Mobile Computer-Adaptive Testing. J Med Internet Res. 2016; 18:e22.

37. Daltroy LH, Liang MH, Fossel AH, Goldberg MJ. The POSNA pediatric musculoskeletal functional health questionnaire: report on reliability, validity, and sensitivity to change. Pediatric Outcomes Instrument Development Group. Pediatric Orthopaedic Society of North America. J Pediatr Orthop. 1998; 18:561–571.

38. Harris PA, Taylor R, Thielke R, Payne J, Gonzalez N, Conde JG. Research electronic data capture (REDCap)-a metadata-driven methodology and workflow process for providing translational research informatics support. J Biomed Inform. 2009; 42:377–381.

39. R: A Language and environment for statistical computing. [http://www.R-project.org/] Accessed 8 April 2018.

40. Surgeons AAoO. AAOS Outcomes Material Users Guide. In:AAOS Outcomes Material Users Guide. Surgeons AAoO. 2001. https://www.aaos.org/uploadedFiles/PreProduction/Quality/About_Quality/outcomes/AAOS%20Outcomes%20Material%20Users%20Guide.pdf. Accessed May 5 2018.

41. Cohen JS, Biesecker BB. Quality of life in rare genetic conditions: a systematic review of the literature. Am J Med Genet A. 2010; 152A:1136–1156.

42. Sajobi TT, Brahmbatt R, Lix LM, Zumbo BD, Sawatzky R. Scoping review of response shift methods: current reporting practices and recommendations. Qual Life Res. 2018; 27:1133– 1146.

43. Van Dijk FS, Sillence DO. Osteogenesis imperfecta: clinical diagnosis, nomenclature and severity assessment. Am J Med Genet A. 2014; 164A:1470–1481.

44. Tournis S, Dede AD. Osteogenesis imperfecta - A clinical update. Metabolism. 2018; 80:27–37.

45. Alcausin MB, Briody J, Pacey V, Ault J, McQuade M, Bridge C, Engelbert RH, Sillence DO, Munns CF. Intravenous pamidronate treatment in children with moderate-to-severe osteogenesis imperfecta started under three years of age. Horm Res Paediatr. 2013; 79:333–340.

46. Balasubramanian M, Parker MJ, Dalton A, Giunta C, Lindert U, Peres LC, Wagner BE, Arundel P, Offiah A, Bishop NJ. Genotype-phenotype study in type V osteogenesis imperfecta. Clin Dysmorphol. 2013; 22:93–101.

47. Bishop N, Adami S, Ahmed SF, Anton J, Arundel P, Burren CP, Devogelaer JP, Hangartner T, Hosszu E, Lane JM, et al. Risedronate in children with osteogenesis imperfecta: a randomised, double-blind, placebo-controlled trial. Lancet. 2013; 382:1424–1432.

48. Andersen PE, Jr., Hauge M. Osteogenesis imperfecta: a genetic, radiological, and epidemiological study. Clin Genet. 1989; 36:250–255.

49. Werfel KL, Hendricks AE. The Relation Between Child Versus Parent Report of Chronic Fatigue and Language/Literacy Skills in School-Age Children with Cochlear Implants. Ear Hear. 2016; 37:216–224.

